# LRT: T Cell Trajectory Inference by Integrative Analysis of Single-Cell TCR-seq and RNA-seq data

**DOI:** 10.1101/2022.04.14.488320

**Authors:** Juan Xie, Gang Xin, Qin Ma, Dongjun Chung

## Abstract

Single-cell RNA sequencing (scRNA-seq) data has been widely used for cell trajectory inference, with the assumption that cells with similar expression profiles share the same differentiation state. However, the inferred trajectory may not reflect true clonal relationships among cells. Single-cell T cell receptor sequencing (scTCR-seq) data provides invaluable insights into the clonal relationship among cells, yet it lacks functional characteristics. Therefore, scRNA-seq and scTCR-seq data complement each other in improving trajectory inference, where a reliable computational tool is still missing. We developed LRT, a computational framework for the integrative analysis of scTCR-seq and scRNA-seq data for T cell trajectory inference. Specifically, LRT utilizes the TCR sequence information to identify clonally related cells and then uses the transcriptomics information from scRNA-seq data to construct clonotype-level cell trajectories. LRT provides a comprehensive analysis workflow, including preprocessing, cell trajectory clustering, pseudotime inference, and marker gene identification. We illustrated its utility using scRNA-seq and scTCR-seq data of CD4^+^ T cells with acute lymphocytic choriomeningitis virus infection, where we could identify cell trajectories that cannot be revealed solely based on scRNA-seq data. Our downstream analyses showed that (i) these trajectories are involved in distinct functional roles; (ii) the expression patterns of their marker genes over the estimated pseudotime nicely coincide with the Th1/Tfh biology that is well established for the CD4^+^ T cell differentiation; and (iii) the higher level of TCR sequence similarities was observed within each cluster, compared to between clusters. The LRT framework was implemented as an R package ‘LRT’, and it is now publicly accessible at https://github.com/JuanXie19/LRT. In addition, it provides two Shiny apps ‘shinyClone’ and ‘shinyClust’ that allow users to interactively explore distributions of clonotypes, conduct repertoire analysis, implement clustering of cell trajectories, and predict cell trajectory cluster marker genes.

**Author Summary:** Understanding the dynamic changes behind biological processes is important for determining molecular mechanisms underlying normal tissue formulation, developmental disorders and pathologies. Usually, a biological process can be characterized by identifying a trajectory, a path that goes through the various cellular states associated with the process. Since cells in different states may express different sets of genes, researchers often infer cell trajectory via capturing transcriptomics changes. Dozens of methods have been developed for cell trajectory inference, and scRNA-seq data is predominantly utilized. However, methods based only on scRNA-seq data cannot tell us if cells from the same trajectory come from the same clone or not. T cells play a key role in the immune system, and their high antigen recognition specificity is largely determined by their TCR sequences. Thanks to the advent of scTCR-seq technology, people can identify the group of cells coming from the same clone. This paper describes our novel computational framework, namely LRT, and demonstrates that by complementing scRNA-seq data with the clonal information from scTCR-seq data using LRT, we are able to identify cell trajectories that cannot be revealed solely based on scRNA-seq data.

## Introduction

Cell trajectory inference aims to understand how cells differentiate, which is a long-standing problem in developmental biology. Understanding how progenitor cells transform into specified functional cells can provide valuable insights into the molecular mechanisms underlying normal tissue formulation, as well as developmental disorders and pathologies [1]. Single-cell RNA-seq (scRNA-seq) data has been widely used to investigate cell trajectories. Assuming that cells in different states express different sets of marker genes, we may order cells along a differentiation trajectory via capturing differentiated transcriptional activities [2]. Based on this rationale, various computational and statistical approaches have been proposed for this purpose, namely trajectory/pseudotime analysis, where well known examples include Monocle [2] and Slingshot [3] (please see Saelens et al. [4] for a comprehensive review). Specifically, Monocle builds a minimum spanning tree (MST) on cells in a low-dimensional independent component space and uses the longest path through the MST as the backbone for pseudotime ordering, while branches were handled by the PQ tree. In the case of Slingshot, in its first stage, it also uses MST to determine the global cell lineage structure. Yet instead of building MST on individual cells, Slingshot constructs an MST on cluster centers, which makes it more robust to noises. Then, in its second stage, it uses a novel method called simultaneous principal curves to obtain smooth branching trajectories. While these approaches have been shown to be useful, they still suffer from intrinsic limitations. Specifically, it assumes that cells with similar expression profiles share similar developmental stages and may come from the same lineage, which might not be always true [1, 5]. Besides, in the absence of clonal information, the inferred trajectory may not reflect the true clonal relationship between cells [1].

The recent advent of single-cell T cell receptor (TCR) sequencing (scTCR-seq) technology has enabled the use of TCR sequences as unique ‘barcodes’ to identify clonally related cells. This is because the cells with identical paired TCR sequences usually arise from the same T cell clone due to the high dimensionality of TCR sequence space [6]. Examining the abundance/proportion of clonal cells and their changes greatly benefits our understanding of adaptive immune response in health and disease [7, 8]; therefore, there has been a rapid accumulation of scTCR-seq data in recent years. Accordingly, several computational and statistical approaches have also been developed, where examples include Immunarch [9] and scRepertoire [10]. However, currently available approaches mostly focus on TCR repertoire analysis, such as clonality, repertoire overlap, repertoire diversity, and gene usage analyses. The utilization of scTCR-seq data for cell trajectory analysis is still limited. In addition, although TCR data provides invaluable insights into the breadth of the antigenic response and clonal relationships between cells, it does not offer information about the functional characteristics of the T cells. Hence, integrating scTCR-seq and scRNA-seq can help elucidate the interplay between TCR sequences and T cell phenotypes. For instance, Zhang et al. [11] observed that TCR similarity constrained the similarity of T cell phenotypes, and TCRs that are more similar to the center TCRs have stronger antigen binding affinity. However, despite such potential to improve T cell trajectory inference, to the best of our knowledge, a computational approach to integrate scRNA-seq and scTCR-seq data for cell trajectory inference is missing in the literature. This is partially due to the statistical and computational challenges related to the high dimensionality of the clonotype space. Specifically, for a given human subject, 10^13^ TCR sequences are typically observed while each clonotype may only span a very limited number of cells. Hence, for effective cell trajectory inference and integration of scTCR-seq and scRNA-seq data, we need a cleverly designed computational framework that can handle these challenges.

In this paper, we propose LRT (**<u>L</u>**ineage inference by integrative analysis of sc**<u>R</u>**NA-seq and sc**<u>T</u>**CR-seq data), a rigorous computational framework for the integrative analysis of scTCR-seq and scRNA-seq data for cell trajectory inference. LRT addresses the complexity and the high dimensionality of scTCR-seq data through a trajectory clustering algorithm and effective information sharing between scTCR-seq and scRNA-seq data. In addition, to support researchers in utilizing the proposed LRT framework for their studies, we developed an R package ‘LRT’, which is now publicly accessible at https://github.com/JuanXie19/LRT. We also developed two Shiny apps, namely ‘shinyClone’ and ‘shinyClust’, and they allow researchers to interactively implement the complete LRT analysis workflow, including exploratory analysis, cell trajectory inference, and marker gene identification. These Shiny apps are provided as part of the R package ‘LRT’.

## Materials and Methods

The workflow of the ‘LRT’ framework is shown in **Fig. 1**, which starts from the integration of scTCR-seq and scRNA-seq data (**Fig. 1A**). Using this integrated dataset, LRT infers cell trajectory by implementing the following four steps. In the first step, it estimates clonotype-level trajectories by building an MST based on the cells sharing the same clonotype (**Fig. 1B**). In the second step, LRT identifies clusters of clonotype-level trajectories using the stochastic block model (SBM) based on the dynamic time warping (DTW) distances (**Fig. 1C**). In the third step, a principal curve is fitted using the cells belonging to the same trajectory cluster, to get the final trajectories and infer the pseudotime (**Fig. 1D**). Finally, differential expression (DE) analysis is implemented to identify genes characterizing each trajectory cluster, namely trajectory cluster markers (**Fig. 1E**).

**Figure 1.**
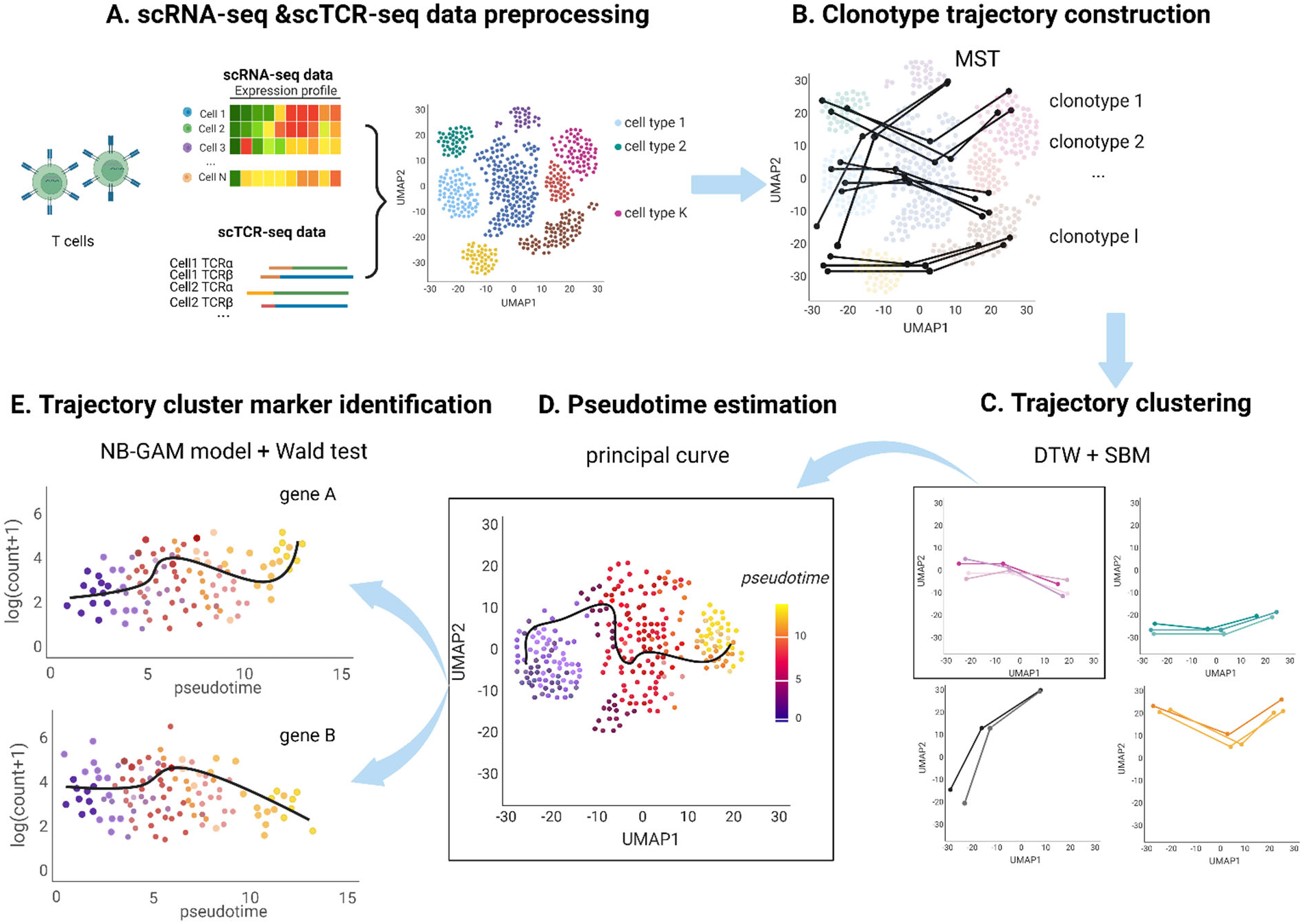
The framework of LRT. **(A)** The preprocessed scRNA-seq and scTCR-seq data are integrated based on cell barcodes. **(B)** Clonotype-level trajectories are first obtained by constructing minimum spanning trees (MST). **(C)** Similar trajectories are grouped using the stochastic block model (SBM), where similarity is evaluated based on the dynamic time warping (DTW) distance. **(D)** For each trajectory cluster, cell pseudotimes are estimated by fitting a principal curve on the Uniform Manifold Approximation and Projection (UMAP) space. **(E)** The relationship between read count and pseudotime is modeled by negative binomial generalized additive model (NB-GAM), and the Wald test is used to determine marker genes.

### Trajectory inference

The core of LRT lies in its trajectory clustering and characterization through the integration of scTCR-seq and scRNA-seq data. It consists of four key steps, including clonotype-level trajectory estimation, trajectory clustering, pseudotime inference, and marker identification (DE analysis). Each step is described in detail below.

### Clonotype-level trajectory estimation

Given the integrated scRNA-seq and scTCR-seq data (see the Section ‘Software implementation’ for more details), LRT infers a trajectory for each clonotype. Specifically, a weighted graph is constructed for each group of cells sharing the same TCR sequence. Here nodes are cell-cluster-specific centers of these cells, while edge weights represent the Euclidean distances between the nodes on the Uniform Manifold Approximation and Projection (UMAP) space [12]. Note that both cell clusters and UMAP in this step are defined based on scRNA-seq data. An MST [13] is then constructed, and trajectories are interpreted as ordered cell clusters by tracing paths through the MST.

### Trajectory clustering

After obtaining the clonotype-level trajectories, LRT conducts a clustering analysis to group similar trajectories, where similarity is defined based on DTW distances [5] between trajectories. DTW distance is a measure to reflect the similarity between two temporal sequences, which is invariant to warping and thus is particularly useful when the two sequences vary in speed. Here we treat each trajectory as time series data, with cell clusters as time points and UMAP coordinates as measurements under each time point. Let’s denote two trajectories *S* = (*s*_1_, *s*_2_,…, *s*_*i*_,…, *s*_*q*_) and *T* = (*t*_1_, *t*_2_,…, *t*_*j*_,…, *t*_*r*_), where *q* and *r* are the numbers of “time points”, respectively. We first build the distance matrix *C* ∈ ℝ^*q*×*r*^ to represent pairwise distances between *S* and *T*, where each element of *C, c*(*i, j*) = ∥*s*_*i*_ − *t*_j_∥, *i* ∈ [1: *q*], *j* ∈ [1: *r*]. Then, we define a warping path as a sequence of points *p* = (*p*_1_,…, *p*_*Q*_) with *p*_*l*_ = (*i*_*l*_, *j*_*l*_) ∈ [1: *q*] × [1: *r*] for *l* ∈ [1: *Q*] to represent a possible alignment between *S* and *T*. The associated total cost/distance for *p* is

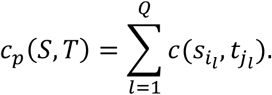

Finally, the DTW distance is defined as the cost associated with the optimal warping path, i.e.,

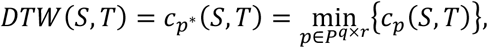

where *P*^*q*×*r*^ is the set of all possible warping paths.

Based on this DTW distance matrix, LRT applies an SBM [14] to identify trajectory clusters (i.e., community detection). SBM is a generative model for random graphs, which is widely employed for recovering (latent) community structures in network/graph data [15]. Let’s denote *n* as the total number of cells. To apply SBM in our case, we convert the DTW distance matrix to an adjacency matrix of a k-nearest neighbor graph, denoted as *A*, where we set 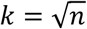 by adopting the widely used heuristics [15, 16]. Given *A* = [*A*_*ij*_] ∈ ℝ^*n*×*n*^ denoting the binary adjacency matrix of a random graph, with elements *A*_*ij*_ indicating the presence or absence of an undirected edge between nodes *i* and *j*, we assume that the absence or presence of an edge between each pair of nodes *i* and *j* follows a Bernoulli distribution, i.e.,

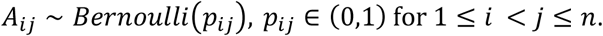

SBM assumes that there is a latent connectivity matrix Θ ∈ ℝ^*b*×*b*^, *b* ≪ *n*, and a map *Z*: {1, …, *n*} → {1, …, *b*}, such that

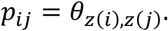

SBM groups *n* nodes into *b* groups, with group labels given by *Z*, and we call these groups communities. A variational Expectation-Maximization algorithm [17] is used for parameter estimation. After fitting SBMs with a different number of clusters, the final number of clusters is determined using the maximum integrated completed likelihood (ICL) criterion [18].

### Pseudotime estimation

As the final step of trajectory inference, LRT obtains the final trajectory by fitting a principal curve [19] on the UMAP coordinates of cells. Let us denote a random vector representing the UMAP coordinates for cells from the same trajectory cluster as *X* ∈ ℝ^2^. In addition, denote *h* as a one-dimensional smooth unit-speed curve in ℝ^2^ parameterized by a single variable *t*. In the principal curve estimation, we iterate the following the projection step and the regularized expectation step:

i. Projection step: Project all cells onto the curve *h*^(*l*−1)^ and calculate the arc length from the beginning of
the curve to each point’s projection, 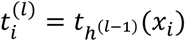, where the projection *t*_*h*_: ℝ^2^ → ℝ^1^ is defined as

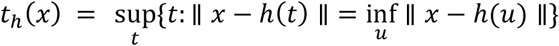
ii. Regularized expectation step: Update the curve *h* by minimizing the distance between data points *x*_*i*_ and those projected onto the curve, 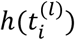.

Here, we interpret the curve *h*(*x*) as the final trajectory for the given cell trajectory cluster and the arc length *t*_*i*_ = *t*_*h*_(*x*_*i*_) as pseudotime for cell *i*.

## Differential expression analysis

While the algorithms described above allow us to infer cell trajectories, it is critical to identify genes characterizing these trajectories to fully understand the biological relevance of these trajectories. To facilitate the interpretation of the inferred trajectories, LRT implements DE analysis for each trajectory cluster. First, for each trajectory cluster, it infers a smooth function of the normalized gene read counts on the pseudotime, using a negative binomial generalized additive model (NB-GAM) [20, 21]. To be specific, for the *v*-th trajectory cluster with *n*_*v*_ cells, let *Y*_*gi*_ be the read counts for gene *g* ∈ {1, …, *G*} and cell *i* ∈ {1, …, *n*_*v*_}, the NB-GAM assumes that

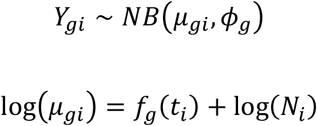

where *µ*_*gi*_ are cell- and gene-specific means, *ϕ*_*g*_ are gene-specific dispersion parameters, *N*_*i*_ is cell-specific offsets that account for sequencing depth, and *f*_*g*_ is the smoothing spline for gene *g*. Here we use a cubic spline function with 6 knots for *f*_*g*_ to balance between the goodness of fit and running time. Finally, a Wald test is used to determine DE genes [21], followed by Bonferroni correction for the multiple testing adjustment.

### Software implementation

The aforementioned LRT framework and the whole analysis workflow were implemented as an R package ‘LRT’ and it is now publicly accessible at https://github.com/JuanXie19/LRT. The LRT software takes both scRNA-seq and scTCR-seq data as input. It accepts scRNA-seq data in the form of a Seurat object [22]. Such integration with Seurat not only allows LRT to be easily integrated into an existing scRNA-seq analysis workflow, but also utilizes Seurat’s powerful features, e.g., correcting batch effects and variations due to technical factors (e.g., sequencing depth) [22, 23]. The Seurat object needs to contain clustering labels and UMAP coordinates for each cell. In the case of scTCR-seq data, it requires a data frame with at least two columns: one column with cell identifiers and another column with TCR sequence information, for which users can provide amino acid or nucleotide sequences of TCR α, β, or paired α-β chain CDR3 region.

### Data preprocessing

LRT integrates scTCR-seq data with scRNA-seq data by attaching the TCR sequence information to the metadata of the Seurat object. LRT also provides some basic quality control functionalities, e.g., checking whether the cell identifiers from the Seurat object match those in the TCR data frame. After the integration, users will get an S4 object containing the updated Seurat object, which is the input for clonotype exploratory analysis and trajectory inference.

### Shiny apps for interactive visualization and analysis: shinyClone and shinyClust

We developed two Shiny apps, *shinyClone*, and *shinyClust*, for interactive and dynamic data exploration and integrative analysis of scRNA-seq and scTCR-seq data.

### shinyClone

*shinyClone* is a Shiny app for exploratory analysis of clonotypes (**Fig. 2**). It provides commonly used repertoire analysis functionalities and lets users intuitively explore how each clonotype is distributed on the reduced dimensional space generated based on scRNA-seq data. *shinyClone* consists of two tabs, one for exploratory analysis of clonotype distribution (tab ‘Clonotype’) (**Fig. 2A**) and the other for repertoire analysis (tab ‘Repertoire’) (**Fig. 2B**). First, on the ‘Clonotype’ tab, users can find a table of clonotypes, along with the relevant information, including (i) how many cells are associated with each clonotype; and (ii) how these cells are distributed among groups (e.g., cell clusters, samples, conditions, or any other groups as available in the metadata of Seurat object) (**Fig. 2A**, left). Users can sort the table by clicking each column name, e.g., to check the most/least abundant clonotypes. By clicking each clonotype in this table, the user can check the distribution of cells associated with each clonotype on the reduced dimensional space, for which the user can choose among UMAP, t-SNE, or principal component analysis (PCA) for the dimension reduction (**Fig. 2A**, middle). A cell density plot is also displayed for clones spanning more than 10 cells (**Fig. 2A**, right). Both plots can be downloaded in the PDF file format by clicking the “Download” button. Second, the ‘Repertoire’ tab provides several common repertoire analysis functions using the R package ‘scRepertoire’ [10], where examples include clonal diversity, clonal homeostasis, clonal proportion, and clonal overlap analyses (**Fig. 2B**).

**Figure 2.**
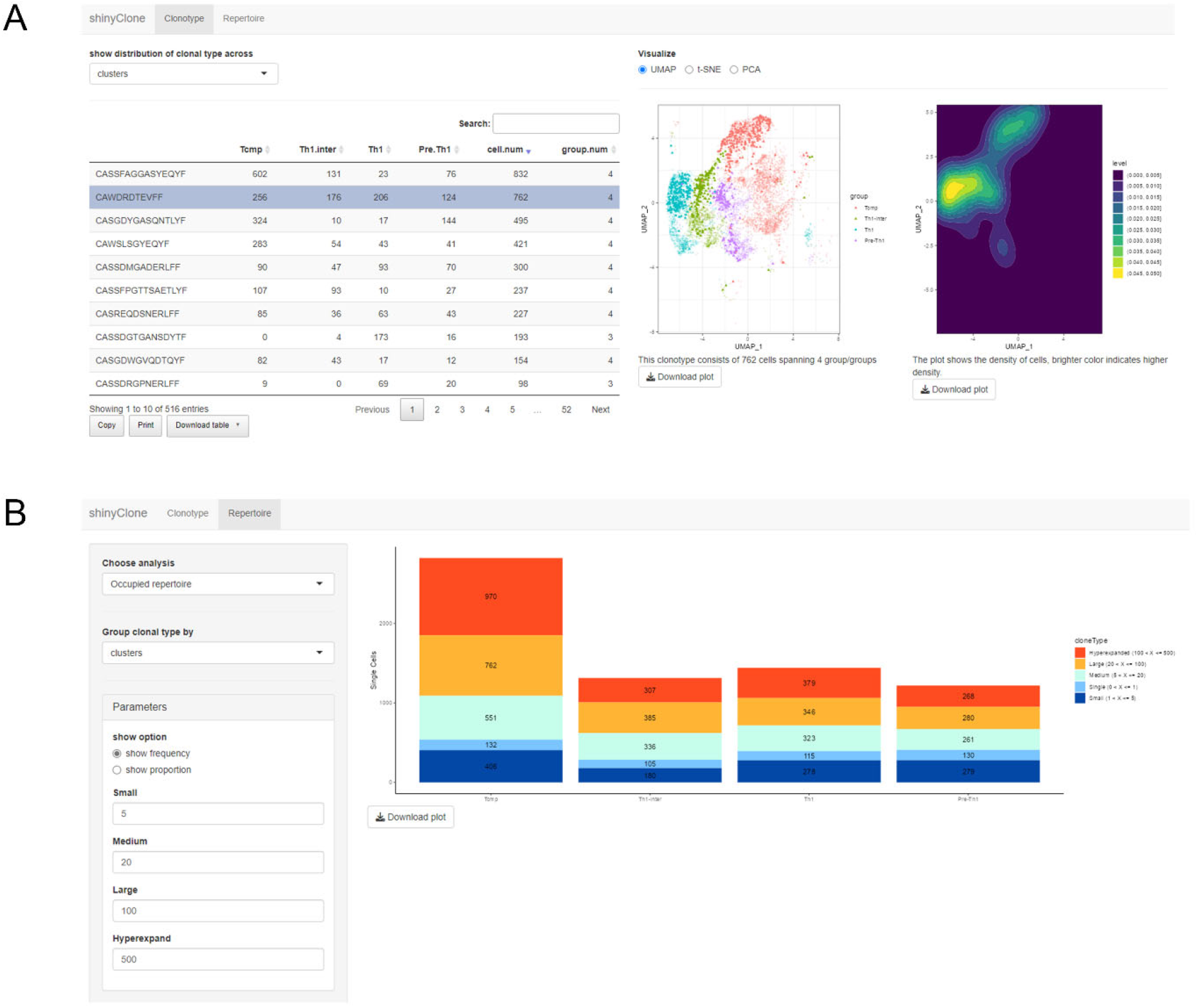
*shinyClone*, a Shiny app for exploratory analysis of clonotypes. **(A)** The ‘Clonotype’ tab allows users to implement various exploratory analyses of clonotype distributions. **(B)** The ‘Repertoire’ tab provides various TCR repertoire analysis functionalities.

### shinyClust

*shinyClust* is another Shiny app that allows users to interactively and dynamically implement the trajectory clustering analysis and marker identification (**Fig. 3**). *shinyClust* consists of three tabs, ‘Clustering exploration’ (**Fig. 3A**), ‘Trajectory clustering’ (**Fig. 3B**), and ‘DE analysis’ (**Fig. 3C**). First, the ‘Clustering exploration’ tab allows users to implement trajectory clustering interactively (**Fig. 3A**). This page shows either ICL or silhouette scores under the current setting, to assist users in evaluating and determining the optimal number of clusters. While SBM is set to be a default here, other clustering algorithms are also provided for flexibility, including hierarchical and partitional clustering. We used the R package ‘sbm’ [24] for the SBM implementation and the R package ‘dtwclust’ [25] for the hierarchical and partitional clustering implementations. Users can also interactively tune parameters for each clustering algorithm. Second, on the ‘Trajectory clustering’ tab, upon choosing a particular cluster to display, *shinyClust* updates (i) a trajectory plot, which shows clonotype trajectories corresponding to the trajectory cluster on the UMAP plot (**Fig. 3B**, the plot on the left, titled ‘Trajectories’); and (ii) a cell distribution plot, which shows the cells associated with this trajectory cluster on the UMAP plot (**Fig. 3B**, the plot on the right, titled ‘Cell distribution’). All the plots can also be downloaded in the PDF file format by clicking the “Download” button. Finally, in the ‘DE analysis’ tab, users can implement DE analysis, which fits the NB-GAM model and conducts the Wald test in the backend using the R package ‘tradeSeq’ [21] (**Fig. 3C**). Here, *shinyClust* shows a table of predicted DE genes with both raw and Bonferroni-adjusted *p*-values, as well as a heatmap of DE gene expression values. If users are interested in how the expression values of a particular gene vary along pseudotime, they can click the corresponding row in the table and the page will show a scatter plot of log read count versus pseudotime, along with the fitted smoothing spline curve.

**Figure 3.**
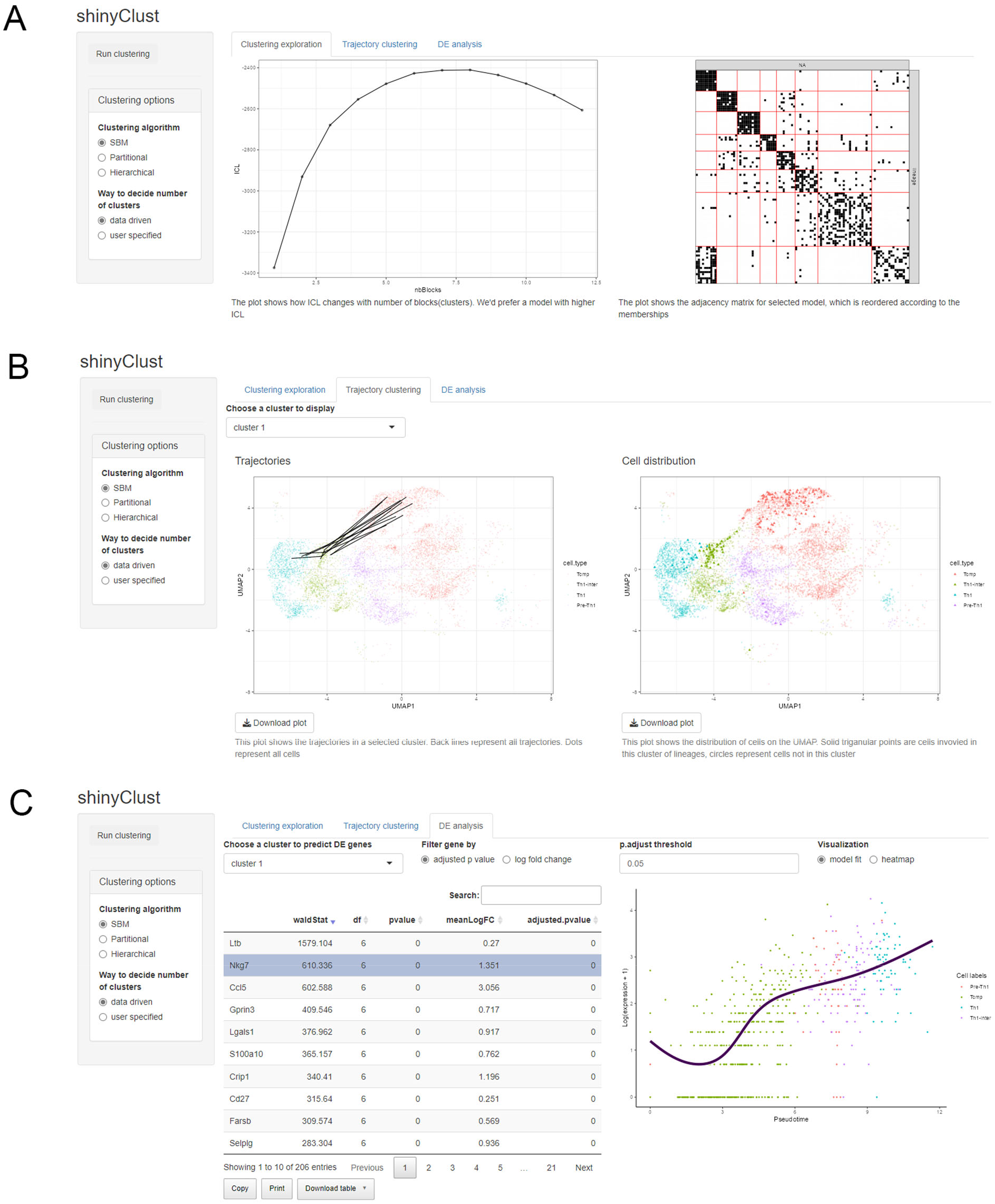
*shinyClust*, a Shiny app for trajectory clustering and marker identification. **(A)** The ‘Clustering exploration’ tab shows the summary of trajectory clustering. **(B)** The ‘Trajectory clustering’ tab shows the trajectory clustering results. **(C)** The ‘DE analysis’ tab allows users to identify marker genes for each trajectory cluster and their changes as a function of the pseudotime.

## Results

### Exploratory analysis of scRNA-seq and scTCR-seq data

To demonstrate the utility of LRT, we analyzed the scTCR-seq and scRNA-seq data generated from the antigen-specific CD4^+^ T cells obtained from five mice during acute lymphocytic choriomeningitis virus infection [26] (GEO accession number: GSE158896). Seurat was used to preprocess scRNA-seq data and implement UMAP dimension reduction and cell clustering. Specifically, for each sample, we filtered out doublets and dead cells (those with a percentage of mitochondrial genes > 10%), calculated cell cycle scores for the remaining cells and regressed out, and normalized using the *SCTransform* approach [23]. Then, the processed datasets from five samples were integrated using the Seurat data integration workflow. Finally, PCA and UMAP dimension reduction were conducted, and then cells were clustered using the Louvain algorithm [27], for which the first 20 principal components and default resolution were used. We obtained 11 cell clusters and annotated them based on the expert knowledge (**Fig. 4**), including T central memory precursor cells (Tcmp) and the cells corresponding to different differentiation stages leading to the type 1 helper T cells (Th1). Specifically, based on the expression of well-known T cell subtype marker genes [26], we annotated cell clusters 0, 1, 4, 9, and 10 as Tcmp, cell clusters 5 and 6 as pre-Th1, cell clusters 2 and 8 as Th1-intermediate (Th1-inter), and cell clusters 3 and 7 as Th1.

**Figure 4.**
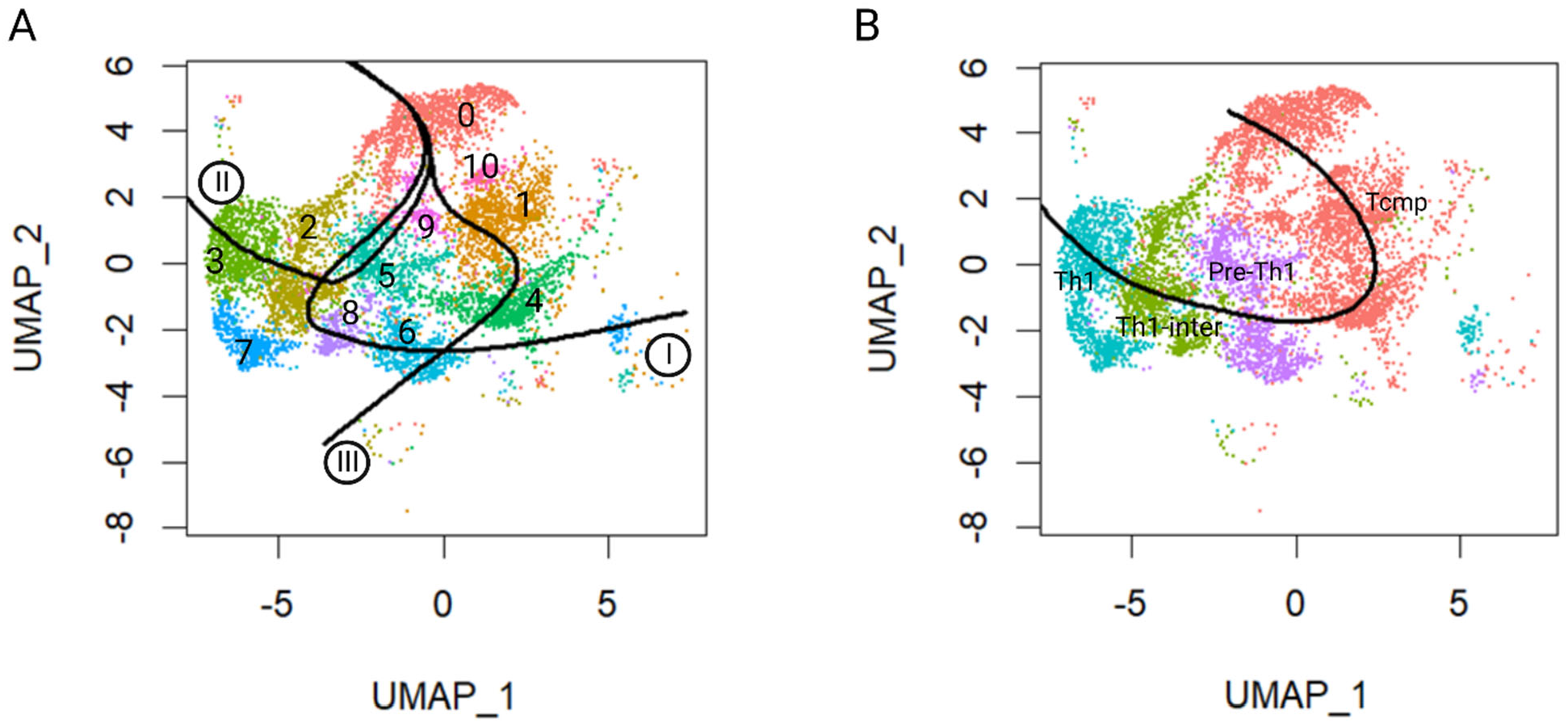
Trajectories inferred based only on scRNA-seq data of the antigen-specific CD4^+^ T cells visualized on the UMAP. **(A)** Cell trajectory inferred based only on scRNA-seq data using the Slingshot algorithm. Dots represent cells and curves represent the inferred trajectories. Here we considered Seurat cell clusters, which are denoted by colors and numbers (0 – 10) in the plot. Roman numbers within circles represent trajectory indices. **(B)** Cell trajectory inferred when we considered the expert-annotated cell types. Cells are colored and labeled accordingly in the plot (salmon for Tcmp, light violet for pre-Th1, green for Th1-inter, and Iris blue for Th1), and the black curve represents the inferred trajectory.

For scTCR-seq data, only the chains that were annotated as full-length and functional were retained. Besides, to avoid any contamination from sorting, only clones having at least two cells from a single sample were considered. After filtering, the TCR sequences from five samples were combined into a single data frame using the R package ‘scRepertoire’ [10]. The resulting scTCR-seq data was then integrated with the above-mentioned Seurat object for the remaining data analyses. Exploratory analysis for the clonotypes was conducted using the *shinyClone* app. There were 516 clonotypes in this dataset and, among those, 69 spanned all four annotated cell types, while 182 of them spanned only one cell type. The top two most abundant clonotypes (defined by CDR3 β chain amino acid sequence) were ‘CASSFAGGASYEQYF’ and ‘CAWDRDTEVFF’, which appeared in 863 and 762 cells, respectively. Most of the cells with the clonotype ‘CASSFAGGASYEQYF’ were Tcmp cells and located at the bottom of the UMAP plot. The cells with the clonotype ‘CAWDRDTEVFF’ evenly spanned all four cell types and these cells mainly occupied the upper part of the UMAP plot.

### Cell trajectory inference

We first inferred cell trajectories based only on scRNA-seq data using the Slingshot algorithm [3] (**Fig. 4**). When we used the Seurat cell clusters (using cluster 0 as the root), Slingshot predicted three trajectories (**Fig. 4A**), including:

- Trajectory I: 0—10—9—5—2—8—7—6
- Trajectory II: 0—10—9—5—2—3
- Trajectory III: 0—10—1—4

When we inferred cell trajectories based only on scRNA-seq data but using four annotated cell types, Slingshot identified a single trajectory of Tcmp – pre-Th1 – Th1-intermediate – Th1 (**Fig. 4B**), which coincides with the known trajectory of Th1 cell differentiation.

Next, we inferred cell trajectories by integrating scRNA-seq and scTCR-seq data using the LRT framework (with SBM as a clustering algorithm). In this case, we discovered five sub-groups of trajectories constituting this trajectory (**Fig. 5A, Fig. S1**). To evaluate whether this trajectory clustering makes sense from the perspective of TCR sequences, we first checked similarities in TCR sequences within vs. between the trajectory clusters. Specifically, we obtained numeric embeddings for TCR sequences using the tessa algorithm [11] and calculated the Euclidean distances between each pair of TCR sequences. We found that within-cluster distances were smaller than or comparable to between-cluster distances (**Fig. 5B**). Among those, within-cluster distances are significantly smaller than between-cluster distances in the case of the trajectory clusters III and IV (p-value=7.38e-14 and 4.01e-7, respectively, one-sided Welch two-sample *t*-test).

**Figure 5.**
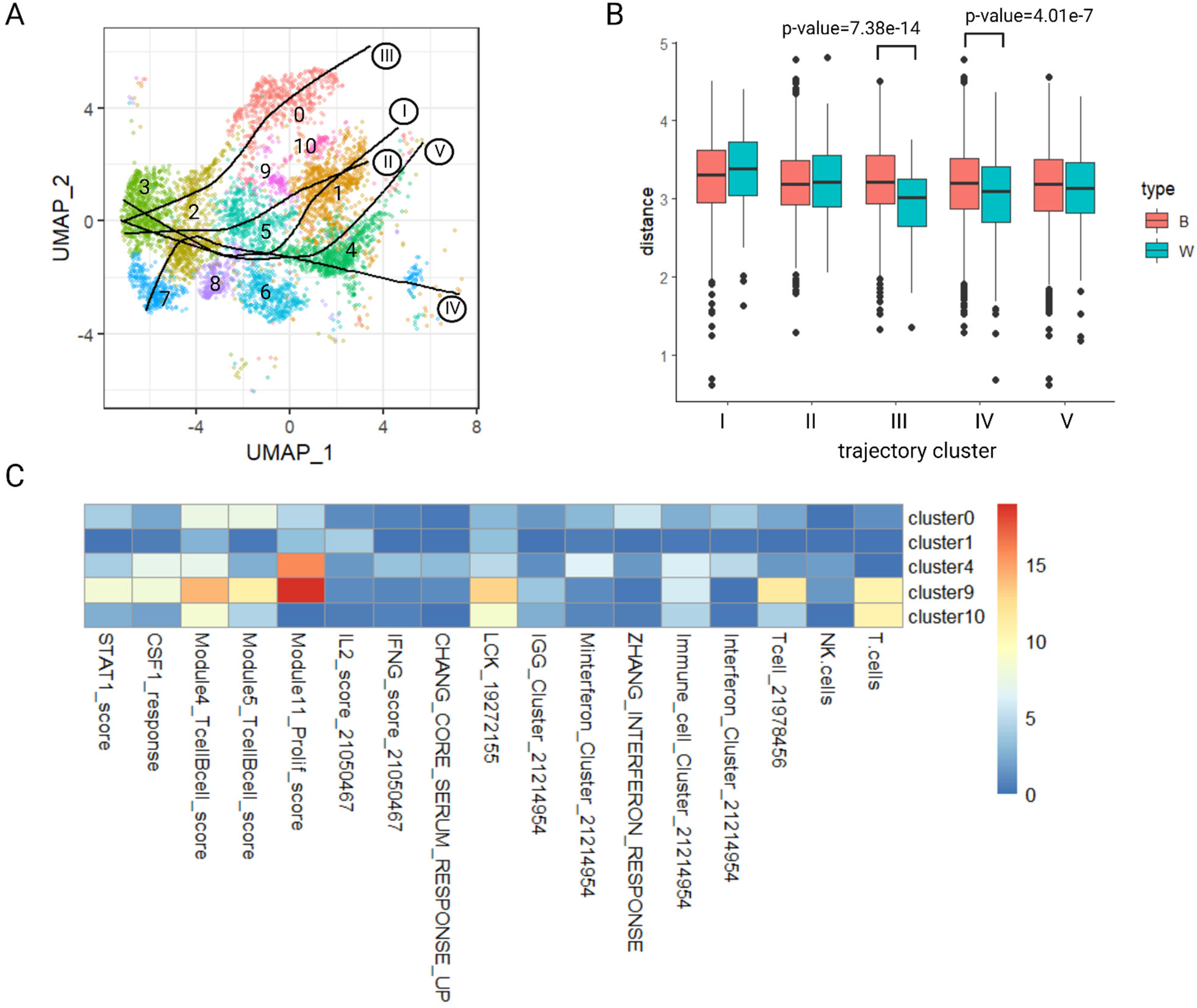
Trajectory inference using the LRT framework. **(A)** Trajectory clusters obtained using LRT. Dots represent cells, while colors and numbers (0 – 10) represent Seurat cell clusters. Each curve indicates the final trajectory for each trajectory cluster, and Roman numbers within circles (I – V) denote trajectory cluster indices. **(B)** Comparison of distances between TCR sequences, where ‘W’ and ‘B’ refer to within- and between-cluster distances, respectively (see the main text for calculation of these distances). **(C)** Enrichment analysis results for the Seurat cell clusters annotated as the Tcmp cell type. The colors represent -log10-transformed adjusted *p*-values.

Interestingly we found that, among the five trajectory clusters, four of them started from the Tcmp cluster but three of them started from different Tcmp sub-clusters. For instance, the trajectory clusters III, I, and V started from Seurat cell clusters 0, 1, and 4, respectively (**Fig. S1**). First, to understand functional differences among them, we identified marker genes for each cell cluster using Seurat and conducted a gene set enrichment analysis using these marker genes. Specifically, hypergeometric tests were applied based on a compilation of 160 immune signatures from the Immune Landscape of Cancer [28], followed by the multiple testing adjustment using the Benjamini-Hochberg procedure [29]. Results indicate that these Tcmp sub-clusters play disparate functional roles (**Fig. 5C**). Specifically, cell cluster 4 was enriched for cell cycle mitotic and cell cycle checkpoints genes, cell cluster 1 was enriched for IL2 activation and signaling pathways, and cell cluster 0 was enriched for immune response and T cell activation. This analysis suggests that integration of scTCR-seq and scRNA-seq data using LRT can reveal the complexity of trajectories that can be missed by the trajectory analysis based only on scRNA-seq data. Next, to understand differences among trajectory clusters from a clonal perspective, we conducted the TCR repertoire analysis based on the Seurat cell clusters. Specifically, we evaluated the overlap of trajectory clusters in terms of clonotype using the Morisita index (**Fig. S2**). Morisita index ranges from 0 to 1 and the larger value indicates a higher extent of overlapping [30]. Repertoire analysis indicates an extremely low overlap of clonotypes among these Tcmp sub-clusters, which further confirmed our clustering results.

Finally, we identified marker genes for each trajectory cluster using the NB-GAM approach (**Table S1**) and evaluated their expressions as a function of the pseudotime. In general, we found that multiple well-established Th1 markers showed increasing expressions while multiple well-known T follicular helper cell (Tfh) markers showed decreasing expressions as a function of pseudotime (**Fig. 6**; **Fig. S3**). For instance, it has been reported that enhanced inhibitor of DNA binding 2 (*Id2*) promotes Th1 cell differentiation while suppressing *Id3* [31, 32]. Therefore, in the process of CD4^+^ T cell differentiating towards Th1 cells, it is expected that *Id2* is upregulated while *Id3* is downregulated, which coincides with our observation for the identified trajectory clusters (**Fig. 6A** and **6B**; **Fig. S3**). Furthermore, chemokine receptor *Cxcr6* serves as a useful marker for Th1 cells [26] while *Cxcr5* is a known marker for Tfh cells. When enhanced *Id2* expression in CD4^+^ T cell shifts the balance of Th1/Tfh towards Th1 cells, *Cxcr6* expression would be up-regulated while *Cxcr5* expression would be suppressed [31]. **Fig. 6C** and **6D** showed patterns in line with this point. Besides the above-mentioned markers, we also observed increasing trends for other Th1-associated markers, such as the Th1 lineage-defining transcription factors *Tbx21, Gzmb, Ly6c2*, and *Cx3cr1*, while observing decreasing trends for the Tfh-specific genes, such as *Ascl2, Il4, Il21*, and *Tox2* [33] (**Fig. S3**). In summary, these expression patterns of Th1- and Tfh-associated markers nicely coincide with the established Th1/Tfh biology, which further confirms the cell trajectories identified using LRT.

**Fig. 6.**
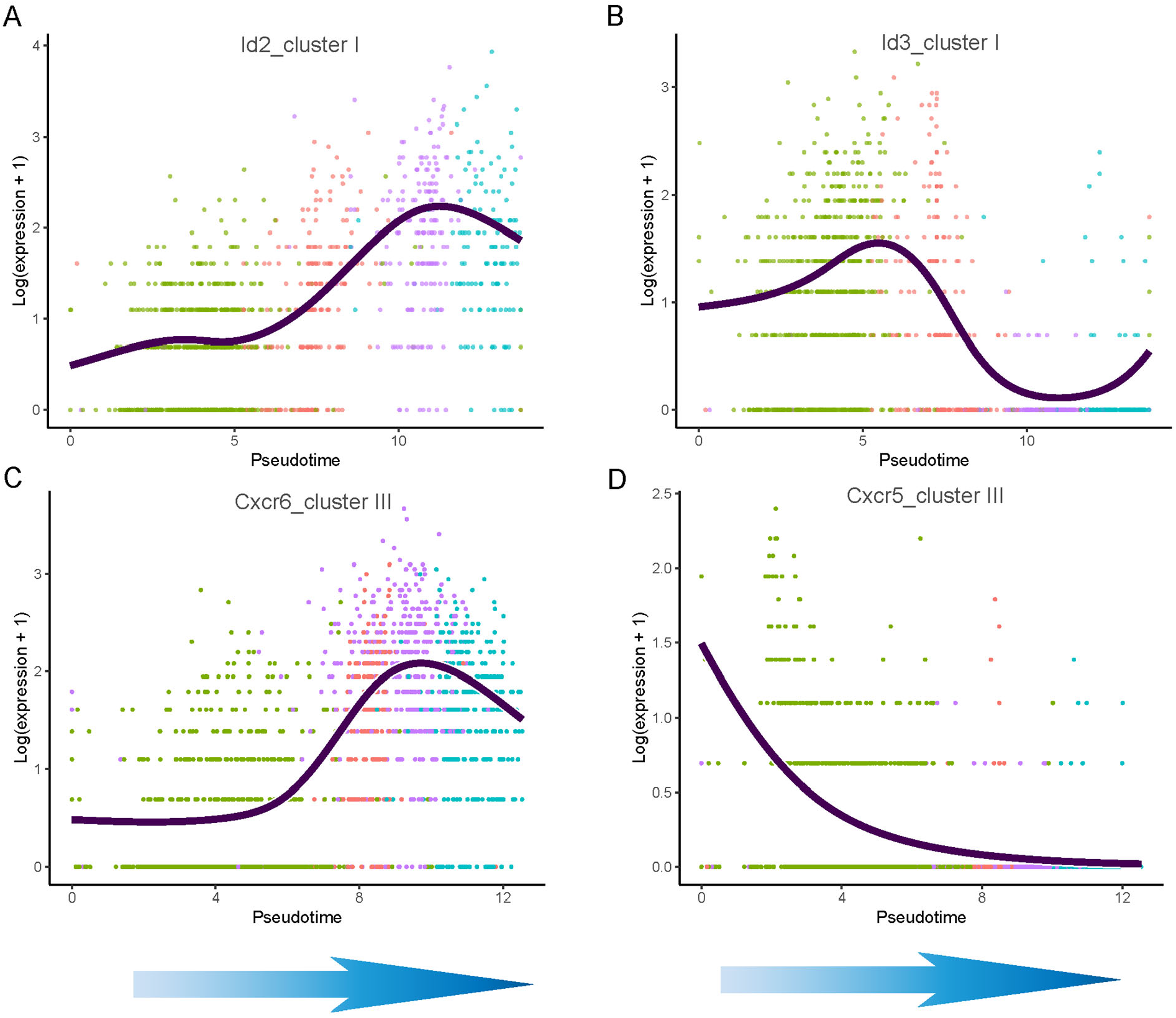
Examples of the trajectory marker genes identified by LRT, which are involved in T cell differentiation. **(A)** and **(B)** Log-transformed read counts of *Id2* and *Id3* genes, respectively, as a function of pseudotime, for the trajectory cluster I. **(C)** and **(D)** Log-transformed read counts of *Cxcr6* and *Cxcr5* genes, respectively, as a function of pseudotime, for the trajectory cluster III. Curves represent smoothing splines fits and dots represent cells. The colors denote different cell types (green for Tcmp, salmon for pre-Th1, light violet for Th1-inter, and Iris blue for Th1). Arrows represent the direction of differentiation.

## Discussion

In this paper, we proposed LRT, a novel computational framework for inferring T cell trajectory by integrating scTCR-seq and scRNA-seq data. LRT addresses the limitation of previous trajectory inference methods that solely rely on scRNA-seq data and neglects the clonal relationship between cells. Specifically, LRT utilizes both the functional information from scRNA-seq data and clonal information from scTCR-seq data. Such integrative analysis allows researchers to identify cell trajectories that cannot be revealed solely based on scRNA-seq data. In addition, LRT provides information about how clonotypes are clustered based on their DTW distances and MST, and this can help researchers decide which clonotypes need to be investigated jointly. Finally, LRT allows researchers to identify marker genes characterizing each of these trajectory clusters through DE analysis based on the principal curve and NB-GAM approaches, and this will help them understand potential drivers of cell differentiation, which is often the ultimate target.

Despite such strengths and power of LRT, it is also not free of limitations, which we plan to investigate in the future. First, while the current LRT framework assumes a single trajectory within each trajectory cluster, biologically various branching events can be possible within each trajectory cluster. We plan to address this limitation by considering more complex trajectory inference algorithms [3, 4]. Second, currently, LRT employs a simple association test evaluating which genes show significant expression changes along the pseudotime. However, researchers might also want to know whether these changes occur more in the earlier stage or in the later stage of the differentiation process. This limitation might be addressed by considering different types of the null hypothesis for the Wald test of NB-GAM. For example, we may modify the contrast matrix so that we can compare gene expression at starting points with those at ending points, as implemented in tradeSeq [21]. Third, currently, LRT implements DE analysis only to determine marker genes characterizing each trajectory cluster. However, researchers might also want to check which genes show DE between trajectory clusters. We plan to investigate approaches like those proposed by Van den Berge et al. [21] to address this. With the aforementioned strengths of the proposed LRT framework and these future development plans, we believe that LRT can be a powerful tool for cell trajectory inference and integrative analysis of scRNA-seq and scTCR-seq data.

## Supporting information

**Figure S1. Clonotype trajectories belonging to each of the trajectory clusters I - V**.

**Figure S2. TCR repertoire overlap between cell clusters**. The numbers within cells show Morisita indices. The numbers on the *x* and *y* axis denote cell cluster indices.

**Figure S3. Additional examples of the trajectory marker genes identified by LRT, which are involved in T cell differentiation**. The left column shows the Th1-associated markers and the right column shows the Tfh-associated markers. Curves represent smoothing splines fits and dots represent cells. The colors denote different cell types (green for Tcmp, salmon for pre-Th1, light violet for Th1-inter, and Iris blue for Th1).

**Table S1. List of DE genes identified using LRT for the antigen-specific CD4**^**+**^ **T cell data**. Each tab provides marker genes for each trajectory cluster. Wald statistics (waldStat), degree of freedom (df), original *p*-value (pvalue), mean log fold change (meanLogFC), and Bonferroni-adjusted *p*-values (p.adj) are reported for each gene.

## Data Availability

The LRT framework was implemented as an R package ‘LRT’ and it is publicly available at https://github.com/JuanXie19/LRT. The Shiny apps ‘shinyClone’ and ‘shinyClust’ are also provided as part of this R package. The data used to produce the results presented in this manuscript is available at GEO with accession number GSE158896.

