## Supplementary figures and images for "LRT: T Cell Trajectory Inference by Integrative Analysis of Single-Cell TCR-seq and RNA-seq data"

### FigS1

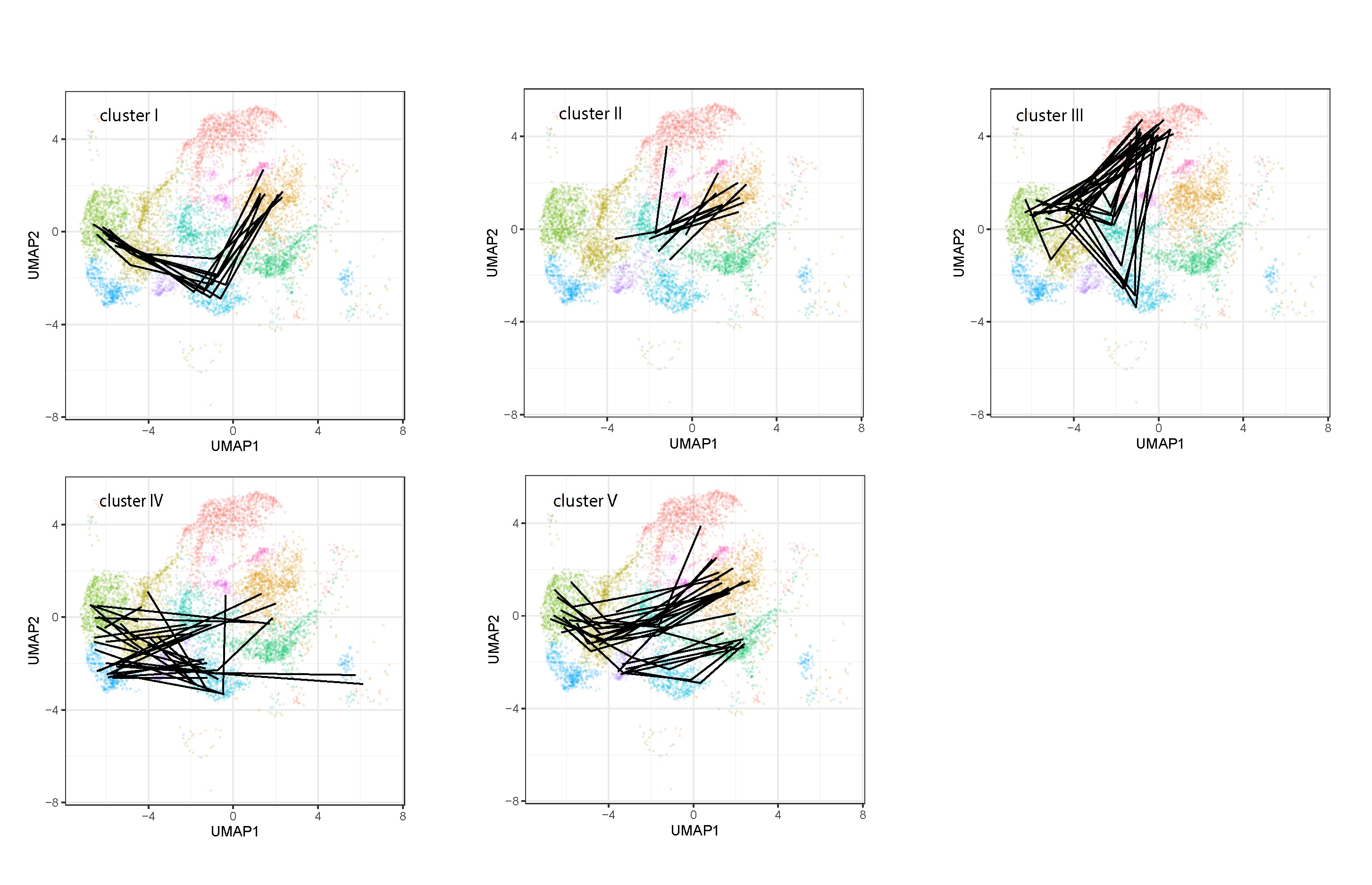

### FigS2

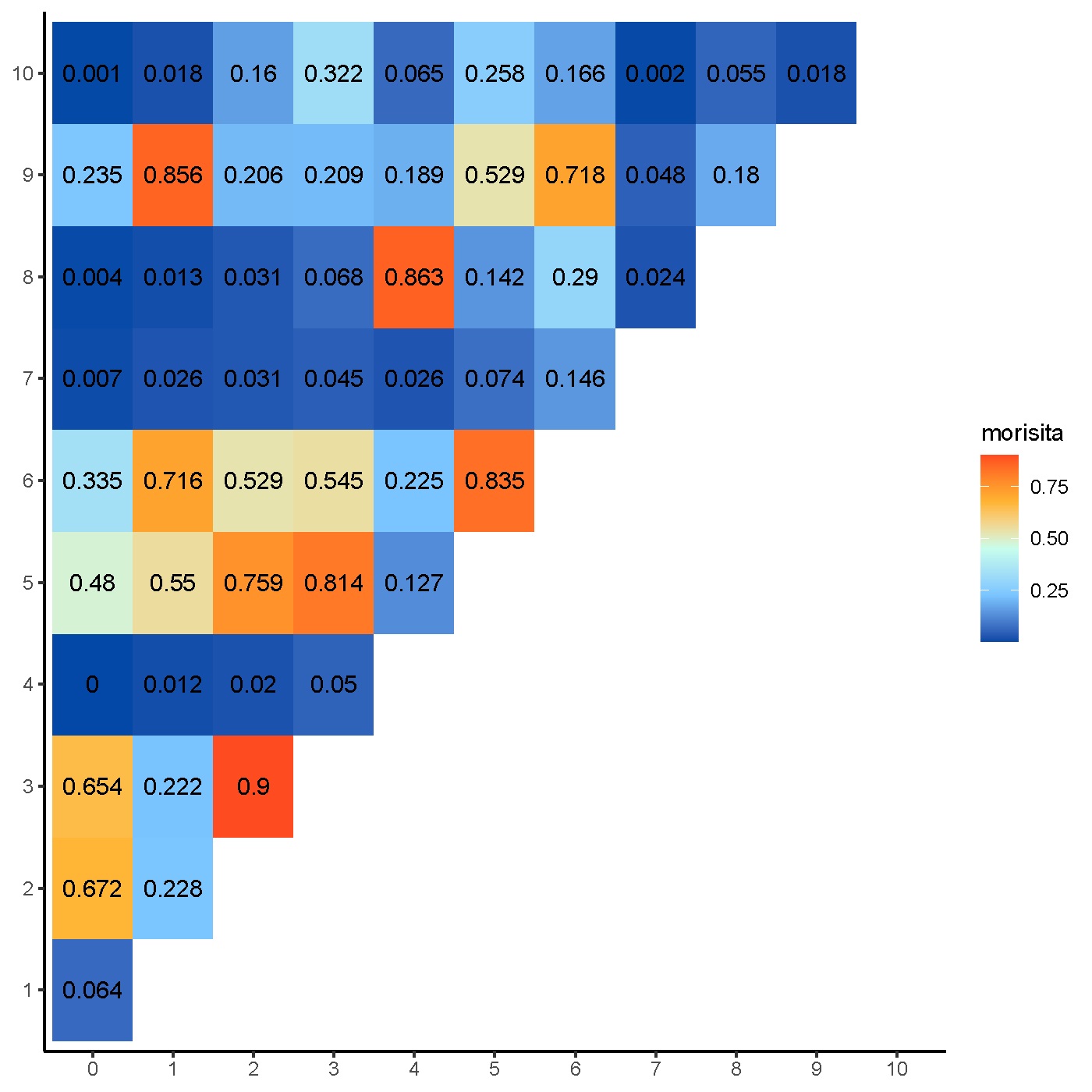

### FigS3

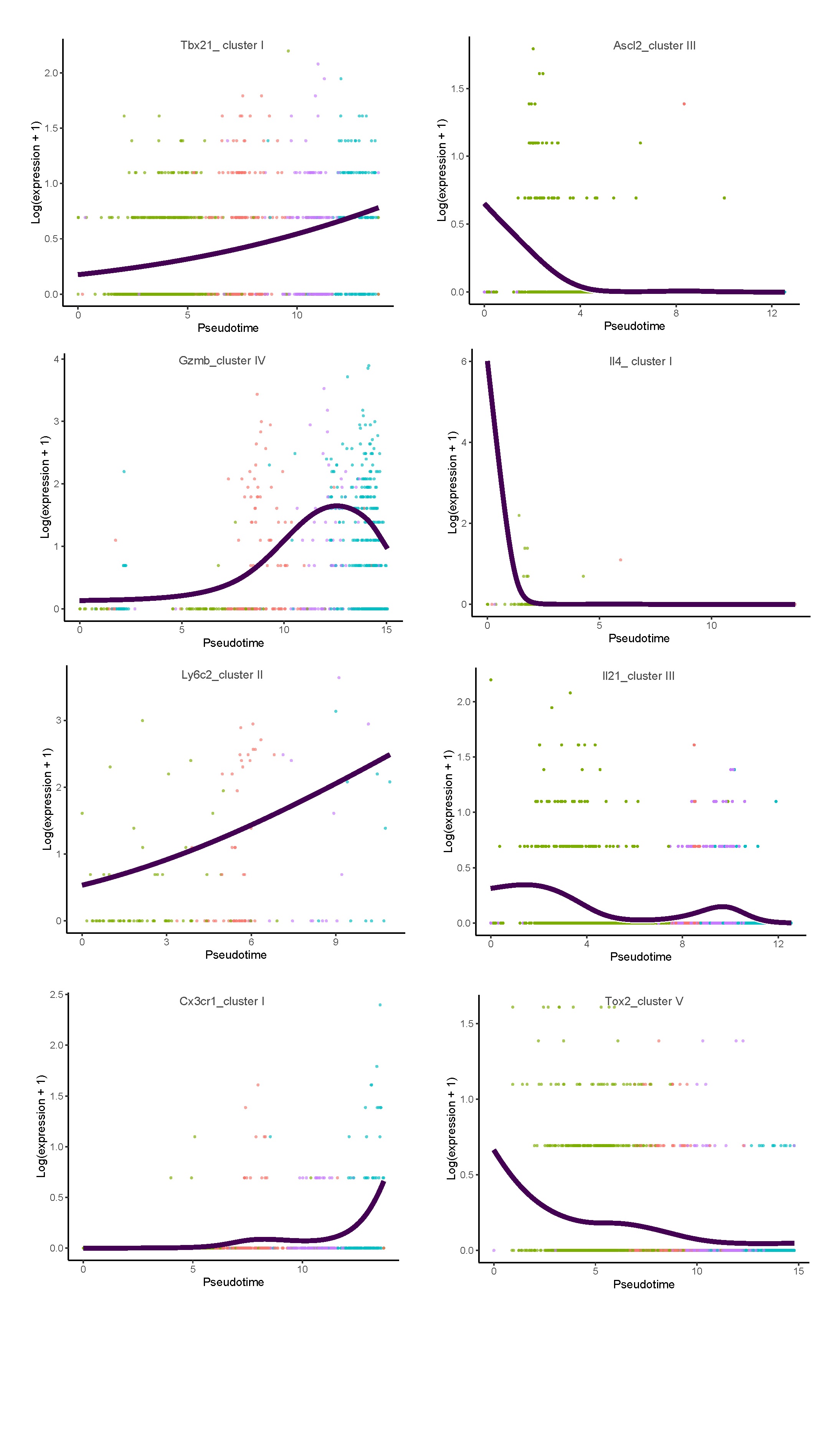
